# Role of miRNA 383 in regulating the mitochondrial machinery and carcinogenesis

**DOI:** 10.1101/2024.04.25.591190

**Authors:** Ashutosh Kumar Maurya, Grace R. Raji, V.B. Sameer Kumar

## Abstract

MicroRNAs are a class of small non-coding RNAs, which regulate the expression pattern of various genes in a mechanism similar to RNAi. Along with their importance in normal physiology, microRNAs play crucial role in cancer initiation and progression. Various microRNAs have been reported to be associated with the important hallmarks of the cancer including altered mitochondrial machinery. With our in-silico analysis we found that miR 383 has targets on crucial mitochondrial genes involved in electron transport chain. So, next we checked the role of miR 383 in modulation of the mitochondrial machinery and effect of this alteration in the process of carcinogenesis. The results suggested that miR 383 contribute to carcinogenesis, possibly by modulating the mitochondrial machinery via targeting ND4L and ATP6 genes involved in the complex 1 & 5 of electron transport chain.

## 1. Introduction

MicroRNAs (miRNAs) are endogenous, small non-coding RNA molecules of approximately 22-25 nucleotides in length (Vaghf A et.al, 2022). They regulate gene expression post-transcriptionally by directly binding to the 3’-UTR of the target mRNAs and exert their function by influencing gene expression through translational inhibition or degradation of target mRNAs (Gulyaeva et.al, 2016, Kittelmann et.al, 2019). In human, endogenous miRNAs regulate almost 30% of genes, thereby playing key role in the regulation of various biological processes that include development, differentiation, proliferation, DNA repair and apoptosis (Menon A et.al, 2022, Arif K. et.al, 2020).

Apart from regulating the normal functioning of the cells, tissue /organ development i.e. heart (Ouyang et.al, 2021), and implications in various disease conditions (Suzuki et.al, 2023), microRNAs also play very crucial role in cancer initiation and progression as they have targets on both the category of genes involved in oncogenesis (Erez Uzunar et.al, 2022), thereby they can regulate the expression pattern of target genes, either at transcriptional or translational level (Liu H et.al., 2012). MicroRNAs can act as tumor suppressor i.e. miR let-7, miR 15/16 family (Otmani K et.al., 2021) or can act as oncogenic microRNA i.e. miR-21 (Ali Sayeda et.al, 2020), which is mainly decided by their down stream impact on the target gene (Wengong et.al, 2019). Apart from carcinogenesis, microRNAs can also regulate the metastasis of cancers (Carla Sole et.al, 2021).

Most important hallmarks observed in variety of cancer includes genetic instability due to the mutations in genome (Yao Y. et.al, 2014), bypassing of anti growth signals, immortality, rapid angiogenesis, metastasis (Hanahan D et.al, 2022) and most importantly the continuous supply of energy to enhance cellular Proliferation (Swapna R et.al, 2022, Nadine D, 2020). In order to achieve the energy requirement, the cancer cells modify the mitochondrial metabolism and activate glycolytic pathway as the alternate source of the energy (Kim S., 2018). Mitochondria are referred as the power house of the cell as it provides energy required by the cellular machinery to complete their actions which includes cell proliferation and differentiation (Osallame et.al, 2012, Spinelli et.al, 2018). As it plays important role in cell growth and division, any malfunctioning in mitochondrial machinery can be directly related to the process of oncogenesis (Patel P et.al, 2022).

Scientific research has established that mitochondrial dysfunction is one of the most prevalent and profound phenotypes of human cancer cells (Wang SF et.al, 2023). Because of their involvement in apoptosis and other facets of tumor biology (Paolina K et.al, 2021), as well as their function as a source of reactive oxygen species, mitochondria have been linked to the carcinogenic process (Narayansami et.al, 2018). Many types of human malignancy such as colorectal, liver, breast, pancreatic, lung, prostate, bladder and skin cancer have been shown to harbour somatic mtDNA mutations (Young Seuok et.al, 2014, Lee H. et. Al, 2009). At organelle level, the micro RNAs having targets on the mitochondrial genes involved in energy production, could play important role in modulating the mitochondrial machinery leading to carcinogenesis (Suriya M et.al, 2022).

Our Insilico analysis revealed that miR 383 has targets on important mitochondrial genes involved in the Electron transport chain of the mitochondria. MiR 383 is located on chromosome 8p22. It has been found to be involved in several cancer types like esophageal squamous carcinoma, hepatocellular carcinoma, ovarian cancer, pancreatic cancer and glioma. We further checked the role of miR 383 in modulation of the mitochondrial machinery and effect of this alteration in the process of carcinogenesis.

## 2. Materials

HeLa, WRL and HepG2 cells were obtained from NCCS (Pune, India); DMEM (Dulbecco’s Modified Eagle Medium), FBS (Fetal Bovine Serum), Antibiotic-antimycotic solution, L-Glutamine, Trypsin EDTA, PBS, Luria Agar and Luria Broth were purchased from HIMEDIA (Mumbai, India); Pro-tease inhibitor cocktail, Actinomycin, Bovine serum albumin (BSA), Gelatin, Sodium dodecyl sulfate (SDS), Ammonium persulfate (APS), Polyethylene glycol (PEG), Formaldehyde, TritonX100, Tween 20, Polyethylenimine (PEI), Sodium Chloride, Acrylamide, Bis-acrylamide, Agarose, Proteinase K, Anti Hexokinase 3 antibody, Anti Histone 3 antibody, Anti CD63 antibdy were obtained from Origin labs (Kerala,India); Beta-actin antibody, Anti-rabbit IgG and Anti-mouse IgG were procured from Sigma–Aldrich (StLuois, MS); PVDF membrane was procured from Merck Millipore (Darmstadt, Germany); Lipofectamine LTX transfection reagent was obtained from Invitrogen (Carls-bad, CA); Anti VDAC antibody was obtained from Santa Cruz Biotechnology (Dallas,TX); MI script II RT kit, QuantiTect Syber Green PCR Mastermix, Universal reverse primer, Plasmid miniprep kit, Plasmid midi prepkit, Gel extraction kit and Quanti Fast Syber Green PCR Mastermix were obtained from Qiagen (Hilden, Germany);

Restriction enzymes, T4 DNA ligase and T4 Polynucleotide Kinase were procured from Thermo Scientific (Waltham, MA); Tris Buffer, Glycine, Methanol and BSA, were obtained from SRL Laboratories (Mumbai, India); Tissue culture plasticwares were procured from NUNC (Roskilde, Denmark); RT-PCR plates and chemilumniscent ECL substrate were obtained from Biorad (Hercules, CA); Trizol and cDNA synthesis kit were obtained from Origin labs (Kerala, India); All the primers were custom designed from IDT (Coralville, IA), Eurofins (Brussels, Belgium) or Xceleris (Gujarat, India); miR 383, MiR 106b, miR 21 and miR 4263 were cloned in the lab by Dr. Grace Raji and Dr. Lincy Edatt; 96 well optiplates were purchased from PerkinElmer (Waltham, MA); pCMVmiR was purchased from OriGene, USA; All glasswares used in the study were procured from Borosil, Mumbai, India; All other plasticwares and chemicals used in the study were from Tarson (Kolkata, India) and Merck (Mumbai, India), respectively, unless otherwise specified.

## 3. Methods

### Cell culture

Hela (cervical cancer cell line), WRL-68 (Human hepatic non cancerous cell line) and HepG2 cells (Hepatic carcinoma cell line) were cultured in DMEM supplemented with 10% FBS, antibiotic-antimycotic solution and L-Glutamine. The cells were maintained under standard culture conditions at 37°C with 5% CO2 and 95% humidity. For experiments, seeding density of 0.4 x 10^4^ cells (96 well), 0.6×10^6^ cells (30mm dish), 0.8×10^6^ cells (60mmdish), 2.2×10^6^ cells (100 mm dish) were used.

### miR 383 cloning

miRNA 383 was cloned in pCMV miR vector between BamH1 and Xho1 restriction sites and successful cloning was confirmed by sequencing.

### Transformation

The competent cells (DH5 α) were transformed with miR 383 plasmid by heat shock method where the plasmid was incubated with the competent cells followed by a quick heat shock at 90 degree C for 2 minutes and then immediately transferring it on ice. The transformed cells were then plated on agar plate containing kanamycin.

### Plasmid isolation

A Single colony was picked from the agar plate and grown in the LB broth containing Kanamycin. The broth was incubated at 37°C in a shaking incubator. The plasmid was isolated by using Himedia midi kit following manufacturer’s protocol.

### Transfection

HeLa cells were seeded in 6 well plates and grown in a monolayer. After reaching 70% confluency, the cells were transfected with miR 383 plasmid using PEI reagent and incubated for 24 hours in a CO_2_ incubator. After 6 hours of the incubation, the medium was replaced with fresh media and further incubated for 24 hours. Following this, the cells transfected with plasmids having fluorescent tags were observed under fluorescent microscope to check the efficiency of transfection

### Isolation of mitochondria

The mitochondria were isolated from the transfected cells using hypotonic buffer, where the cells were allowed to swell in the buffer for 10 minutes and then break open the cells to release the mitochondria. The cell suspension was then centrifuged at 1300g to remove the cell debris, followed by centrifugation at 12000g to get the mitochondrial pellet. The mitochondrial pellet was suspended in the mitochondrial resuspension buffer.

### Sonication

The mitochondrial pellet was mixed with Lysis buffer and sonicated for 2 minutes at 70% amplitude with 15 sec ON and 30 sec OFF cycle on 4°C. The solution obtained, was centrifuged at 12000g for 10 minutes. The supernatant was collected and protein estimation was done followed by sample preparation for SDS PAGE.

### Protein estimation

Protein level of mitochondria was estimated by Bradford method (Bradford etal,1976). To achieve this, 10μl of sample and 90μl of bradford reagent (50 mg Coomasie Brilliant Blue-G250 in 25ml ethanol and 50ml of phosphoric acid made upto 100ml with water) was added in triplicates in 96 well plate and the absorbance was taken at 595nm by multimode plate reader. The concentration of protein was calculated from the standard plot to BSA with concentration range from 10μg-100μg.

### SDS-PAGE

Protein sample was prepared by mixing of 6x SDS loading dye and boiling it at 90°C for 10 minutes in water bath. The sample was immediately kept on ice and briefly centrifuged before loading on SDS-PAGE gel. The electrophoresis was carried out by using Bio-Rad electrophoresis unit. The protein samples were run through the stacking gel at 80V for 15 minutes and through the resolving gel at 100V at room temperature until the dye reached the end of the gel.

### Western blot analysis

The purity of the mitochondrial pellet was checked by western blot using mitochondria specific antibody (VDAC). Also, the mitochondrial pellet was checked for the nuclear and cytoplasmic contaminants using Histone H3 antibody for Nucleus and Hexokinase HK3 antibody for the cytoplasm.

### RNA Isolation

RNA was isolated from the mitochondrial pellet as well as from the total cell using trizole reagent. Following this, the concentration of the RNA was checked by using nano drop.

### Polyadenylation of RNA

Poly A tail was added to the RNA by Poly A Polymerase enzyme, using manufacturers protocol. This reaction set up was incubated at 37°C for 30 minutes followed by heat inactivation for 5 minutes at 65°C.

### cDNA synthesis

The polyadenylated RNA were used for the synthesis of miRNA 383 specific cDNA by Kang method. Apart from this, total RNA was used to synthesize the cDNA for checking the expression of mitochondrial genes.

### Real Time PCR (qRT PCR)

Quantative real time PCR was performed to check the expression pattern of the microRNA 383 and other mitochondrial genes in mitochondria before and after over expression of miR 383.

### mRNA stability assay

To elucidate the targeting of mitochondrial genes by miR 383, mRNA stability assay was performed. The cells were transfected with miR 383 using PEI method. 24 hours post transfection the cells were treated with actinomycine D at 0, 1, 3, 6 and 12 hours. The samples were collected at each time point for gene expression study.

### Oxygraph analysis

To check the phenotypic effects of the down regulation of the mitochondrial genes by miR 383, the oxygraph analysis was performed, where the oxygen consumption level was checked in the miR 383 over expressed samples and compared with the control samples. In brief, the cells were grown in a 6 well plate and transfected with the candidate microRNAs using PEI method. After 48 hours of incubation at 37°C, the cells were trypsinized and the cell pellet was resuspended in respiration buffer. Later, 1ml of the cellular suspension was added to oxymeter and oxygen intake reading was recorded for 10 minutes. The readings were used to plot the graph to represent the oxygen consumption by the mitochondria.

### Exosome isolation

The exosomes were isolated from the miR 383 over expressing cells using PEG method. To achieve this the cells were transfected with miR 383 and media was replaced with serum free medium. 48 hours post transfection; the spent media was collected and centrifuged at 2000g for 1/2 hour, to remove cell debris. Following this, the cells were mixed with PEG solution in 1:2 ratios and incubated at 4 degree C overnight. Finally, the solution was centrifuged at 12000g for 1 hour. The exosomal pellet was dissolved in PBS and protein estimation was done.

### Cell migration Assay

Cancer cells have the metastatic properties, where they move from its origin to another place and form a secondary tumor. To mimic this in-vitro we perform the cell migration assay with an objective to check, if the exosomes from miR 383 over expressing cancer cells could induce cellular migration of normal Hepatic cells. A scratch was made in the WRL monolayer and treated with exosomes isolated from miR 383 over expressing cancer cells. The cells were allowed to fill the gap formed by the scratch for 48 hours and the images were taken 0, 24, 48 hours respectively and quantified using ImageJ.

### Soft Agar Colony formation assay

To elucidate the role of miR 383 in inducing carcinogenic properties in normal heaptic cells, the colony formation assay was performed where the WRL cells were suspended in the low melting agarose and treated with the exosomes. The cells were incubated at 37°C for 28 days and allowed to form the colonies. The colonies were stained with Coomasie brilliant blue and images were taken from 20 different locations and quantification was done by using ImageJ.

### In-Silico analysis

miR 383 target prediction analysis and scoring for mitochondrial genes was done using five algorithms i.e. MirBase, miRanda, Target Scan, miRDB and MicroRNA.org. Five highest scored mitochondrial genes (ND6, ATP6, Cyto-B, Cox1, ND4L) were selected for target validation.

### Statistical analysis

All the data in the study were expressed as the mean with the standard error mean of at least three experiments, each done in triplicates. SPSS 11.0 software was used for analysis of statistical significance of difference by Duncan’s One way Analysis of Variance (ANOVA). A value of P<0.05 was considered significant.

## 4. Results

### Differential levels of miR 383 in the mitochondria of hepatic cancer cell (HepG2) and non-cancerous hepatic cell (WRL)

As first part of the study, we checked the microRNA 383 enrichment in HepG2 and non-cancerous hepatic cell line (WRL) by qRT-PCR. The results revealed that, there was negligible difference in the expression of miR 383 in HCC and normal hepatic cells. Following this, we checked the levels of miR 383 in the mitochondria of cancerous & non-cancerous hepatic cells and the results suggested that the abundance of miR 383 was 3.5 fold lower in the mitochondria of the HepG2 cells when compared with the control cells (WRL).

**(Figure 4.1-Supplementary data)**

### miR 383 gets targeted to the mitochondria in a selective manner

As the microRNA 383 was found significantly low in the mitochondria of HepG2 cells, we over expressed miR383 in HepG2 cells and checked its relative levels in the cytoplasm & mitochondria. The results of experiment suggested that miR 383 gets targeted to the mitochondria in a preferential manner, with a 6 fold increase in the levels of miR383 in the mitochondria and 2 fold increase in the cytoplasm when compared to the levels in mock transfected cells.

**(Figure 4.2-Supplementary data)**

### MicroRNA 383 down regulated all the candidate mitochondrial genes

Since, the target prediction and scoring analysis of miRNA 383 revealed that it has target on important mitochondrial genes. So, after confirmation of the preferential localization of miR 383 in the mitochondria, next we checked the effect of the higher levels of miR383 on the expression pattern of their target genes in the mitochondria. The results suggested that the levels of all the miR 383 target genes were significantly lowered, confirming the in-silico target prediction analysis. It was found that ND4L gene was 2 fold lowered and the levels of ATP6, Cyto B, ND6 and Cox1 were found to be 1.8, 1.6, 1.9 and 1.7 fold lowered respectively.

**(Figure 3-Supplementary data)**

### 3’ UTR analysis and mRNA stability assay confirmed the targeting of mitochondrial genes

With the target validation study, we found that the expression levels of all the five candidate mitochondrial genes got down regulated by miR 383. To check if these genes are directly targeted by miR383, we performed mRNA stability assay and the results revealed that the levels of ATP6 and ND4L mRNA was significantly reduced under condition of miR 383 over expression. However, a lowering pattern, though not significant was observed for the levels of Cox1, Cyto B & ND6 mRNA under miR 383 over expressed condition, suggesting that out of 5 target genes, ATP6 & ND4L are effectively targeted by miR 383 and falls in line with the 3’ UTR analysis, suggesting that 3’ UTR of these genes harbor putative miR 383 binding sites.

**(Figure 4.4-Supplementary data)**

### Over expression of miRNA 383 reduces the oxygen consumption

With the results obtained from mRNA stability assay, we found that miR 383 effectively targets ATP6 and ND4L gene, which play very crucial role in the electron transport chain of the mitochondria and any change in the expression pattern of these genes would significantly alter the normal functioning of the mitochondrial system. To elucidate the phenotypic impact of this down regulation on the functioning of the mitochondria, we perform the oxygraph analysis by checking the levels of the oxygen consumption by the cells. The result of the oxygraph analysis revealed that the levels of oxygen consumption by the cells went significantly down under miR 383 over expressing condition when compared with the control cells, suggesting its role in dysegulating the mitochondrial machinery by targeting ATP6 and ND4L genes, involved in complex 1 and 5 of electron transport chain of the mitochondria.

**(Figure 4.5-Supplementary data)**

### MicroRNA 383 escalates the rate of cellular migration in normal as well as cancerous hepatic cell lines

Our previous results of target validation and mRNA stability assay revealed that miR 383 gets targeted to the mitochondria and down regulated the expression of the ND4L & ATP6, important in ETC, and it resulted in the reduced oxygen consumption by the HepG2 cells, possibly due to the altered mitochondrial machinery. So, we next checked the role of this altered mitochondrial metabolism in carcinogenesis.

The results of cell migration assay revealed an increase in the rate of cellular migration when treated with exosomes isolated from the miRNA 383 over expressing cells when compared to the cells treated with the exosomes isolated from the mock transfected cells and the pattern was found to be persistent in both, cancerous as well as non-cancerous hepatic cell lines.

**(Figure 4.6 & 4.7 -Supplementary data)**

### miR 383 enhances colonogenic property of cancerous and non-cancerous hepatic cell lines

Along with cell migration assay, we also performed the colony formation assay to check the carcinogenic ability of miR 383 in normal and cancerous hepatic cells. Colony formation assay result shown an increase in the number and size of the colonies when treated with the exosomes enriched with miR383 when compared with the colonies treated with exosomes isolated from the mock transfected cells.

**(Figure 4.8-Supplementary data)**

## 5. Discussion

MicroRNAs are a class of short non-coding RNA (Vaghf A. et.al, 2022) that interacts with the 3’UTR of mRNA to disrupt protein synthesis, thereby controlling the expression of its target genes (Zhang W et.al, 2013, Tufecki et.al, 2014). MicroRNA is known to play a crucial role in the development of a number of diseases (Konovalova et.al, 2019), including heart disease, neurological disorders, cancer and others (Maqbool R et.al, 2014, Chenyu J et.al, 2019), in addition to regulating genes involved in normal cellular activities (Guz M et.al, 2014). miRNAs act as the key players in the initiation and progression of cancers (Yang G et.al, 2014). Via their involvement in a variety of biological processes such as cell proliferation, regeneration, differentiation etc., microRNAs may either promote or hinder a disease scenario (Gwad et.al., 2018, Qie S et.al, 2016).

MicroRNA-383 is located on P arm of chromosome 8, within the third intron of the sarcoglycan zeta (SGCZ) gene (Yi Q et.al, 2022). Micro RNA-383 falls in the category of tumor suppressive microRNAs which has been found to be downregulated and plays a suppressive role in a variety of cancers, including glioma, breast cancer etc. (Bartel D et.al, 2009, Jianchau et.al, 2019). Hepatocellular carcinoma (HCC) is the third fatal cancer, accounting for major cancer related deaths worldwide (Yang JD et.al, 2010, Sieno S. et.al, 2018). It is the prevailing histological type in liver cancer, with yearly 3% increase in the total number of cases (Siegal et.al, 2017). miR 383 expression has been reported to be significantly high in the advance stage of HCC (as revealed by the transcriptome data analysis), in contrary to that some studies has also suggested that miR 383 inhibits tumor growth and initiates apoptosis in HCC. (Fang et.al, 2017).

One of the major hallmark of cancer is dysregulated mitochondrial machinery (Monsalve et.al, 2007, Sripada et.al, 2012), where cancer cells modulate the mitochondrial metabolism to generate the continuous supply of energy. The cancer cells alter the ETC and activate the glycolytic pathway as alternate pathway of energy needed for their exponential growth. MicroRNAs could play important role in this switching of ETC to glycolysis, as they have targets on important mitochondrial genes involved in the normal functioning of the mitochondria. This class of microRNAs are called mitomiRs, which can be either transcribed from nuclear genome or mitochondrial genome and localized in mitochondria (Mora Morri et.al, 2017). MitomiRs are considered as the primary regulators of mitochondrial metabolism (Song R et.al, 2019).

The primary objective of this study was to elucidate the role of miR 383 in modulating the mitochondrial metabolism and effects of this reprogramming, if any, on oncogenesis. Since, miR 383 is a nuclear coded microRNA, first, we checked its presence in the mitochondria by qRT PCR analysis and found that its expression was very low in the mitochondria. Recent studies suggest that nuclear coded micro RNAs gets targeted to the mitochondria (Huaping Li et.al, 201). So we over expressed miR 383 in HepG2 cells and the qRT PCR data revealed that, miR 383 got enriched in the mitochondria by 20 folds, in a preferential manner.

It was seen that the levels of miR 383 was significantly low in the mitochondria of HepG2 cells. Further it was also seen that when over expressed, miR 383 was preferentially localized to the mitochondria. Also, we have seen in the insilico analysis that in majority of HCC, the levels of miR 383 is significantly higher. In this context, increasing the levels of miR383 in HepG2 cells (which otherwise don’t show any difference) and its targeting to the mitochondria provided us with an apt model to understand the mitochondrial role of miR 383 in HCC.

The In-Silico analysis, revealed that miR 383 has targets on 5 important mitochondrial genes i.e., ATP6, ND4L, ND6, Cyto B and Cox1, crucial for complex 1, 3, 4 and 5 of the electron transport chain (ETC). To check it further, we over expressed miR 383 and performed a target validation study to check whether miR 383 could get targeted to the mitochondria and alter the expression of its mitochondrial target genes. The qRT PCR results of miRNA 383 over expression, shown a selective enrichment of the miR 383 in the mitochondria and the target validation experiment suggested that miR 383 significantly lowered all of its target genes, most significantly, ATP6 and ND4L. Next, we performed mRNA stability assay, to further prove the targeting of ATP6 and ND4L by miR383. The results revealed that ATP6 and ND4L gets targeted by miR 383 and falls in line with the our 3’ UTR analysis that revealed that 3’ UTR of ATP6 and ND4L harbour putative miR21 binding site. Along with ATP6 and ND4L, we found that Cox1, Cyto-B and ND6 also gets targeted by miR 383, however, to a lesser extent.

Various studies have shown that overexpression of miR-383 can regulate the mitochondrial functioning (Fang et.al, 2017). This regulation could be due to the suppression of the expression of the mitochondrial genes.

In this study we found that ND4L and ATP6 were direct target of miR 383 and these genes are crucial for the mitochondrial machinery. So, the down regulation of these genes would directly contribute to the dysregulated mitochondrial metabolism. To check the phenotypic effect of this down regulation, we performed oxygraph analysis, where we measured the oxygen consumption by mitochondria of miR 383 over expressing cells. The oxygraph analysis shown a decrease in the oxygen consumption by miR 383 over expressing cells, suggesting that miR 383 possibly altered the mitochondrial machinery by down regulating the expression of the two important genes i.e.ATP6 and ND4L, involved in the complex 1 and complex 5 of the ETC. Several reports have demonstrated that miR-383 mediates cellular proliferation via regulating the expression pattern of proliferation-associated genes i.e. Cyclin D1 (Yi et.al, 2022). Cyclin D1 is well known in regulating cell proliferation and has been demonstrated to be over expressed in several cancers (Qie S et.al, 2016). Based on the fact that MiR 383 alters the cell proliferation, next we checked if miR 383 could induce the carcinogenesis in hepatic cell lines

To achieve this, we performed cell migration assay and soft agar colony formation assay by treating the hepatic cell line with the exosomes enriched with miR 383. The results of the cell migration and colony formation assay suggested that migratory and colony formation ability of the cells got enhanced when treated with the exosomes isolated from miR 383 over expressing cells. Therefore, all these results together, suggests that miR383 possess carcinogenic ability and it possibly involve the modulation of mitochondrial metabolism.

## Acknowledgment

We acknowledge Indian Council of Medical Research, Ministry of Health, Govt. of India for the financial assistance in form SRF and Kerala state council for Science Technology & Environment, Govt. of Kerala for fellowship in the form of JRF and SRF to Mr. Ashutosh K. Maurya. We also acknowledge Central University of Kerala for providing all the necessary facilities to carry out this research work.

## Author Contributions

The authors confirm contribution to the paper as follows: Study conception and design: VBSK, Bioinformatics and wet lab work: AKM. Cloning of miR 383: GRR. All authors reviewed the results and approved the final version of the manuscript.

## Supplementary Data

**Figure 4.1:**
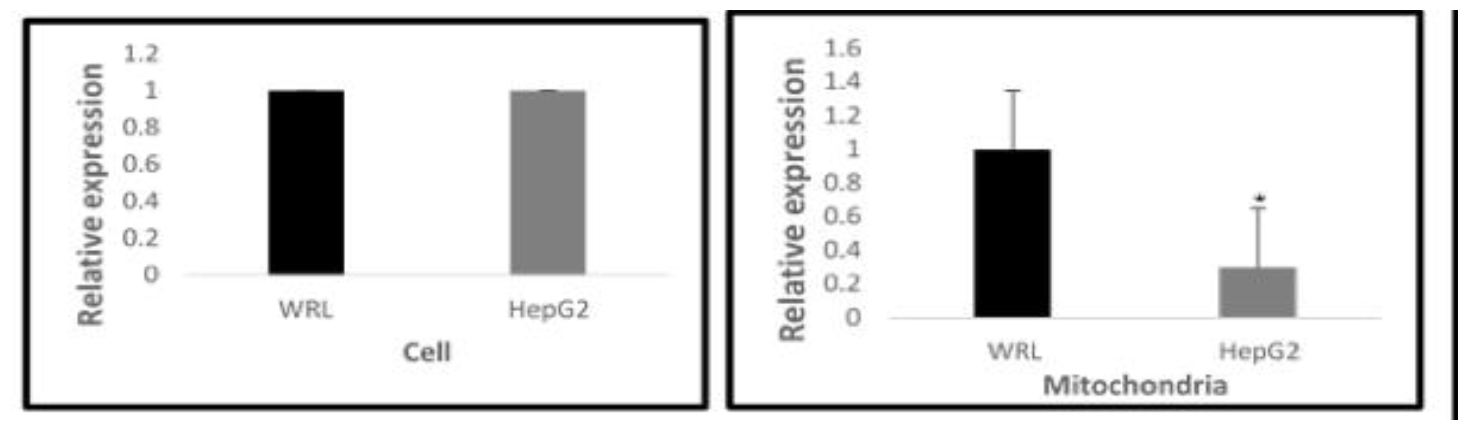
Differential levels of miR 383 in cytoplasm and mitochondria. RT PCR analysis was performed to check the expression pattern of miR 383 in mitochondrial and cellular fraction of HepG2 cells, keeping WRL as control. The results suggested that levels of miR 383 was significantly low in the mitochondria of HepO2 cells. A.) Relative expression levels of miR 383 in the cells B.) Relative levels of miR 383 in the mitochondria. Results presented are average of three experiments ± SEM each done at least in triplicate, p<0.05. .* ‘Statistically significant when compared to control.

**Figure 4.2:**
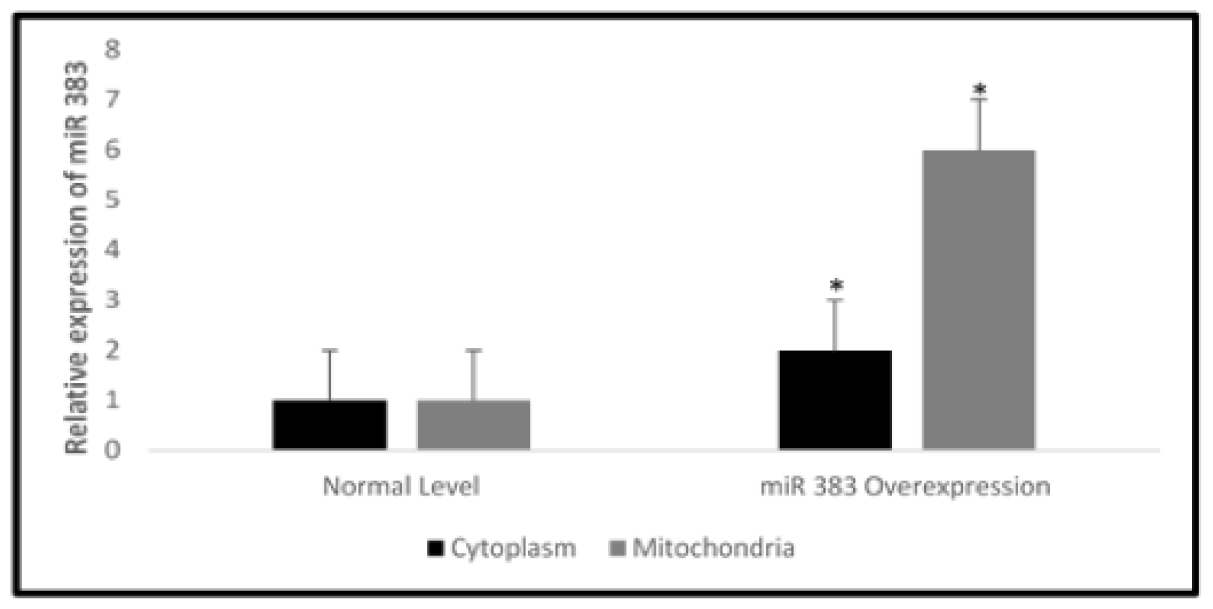
miR 383 gets targeted to the mitochondria in a selective manner. miR 383 was over expressed in HepG2 cells followed by purification of mitochondria & isolation of RNA from mitochondrial as well cis cytoplasmic fractions and, qRT PCR was then performed to check the levels of miR 383. The results revealed that miR 383 gets targeted to mitochondria in a selective manner. Results presented are average of three experiments + SEM each done at least in triplicate, p<0.05.*Statistically significant when compared to control.

**Figure 4.3:**
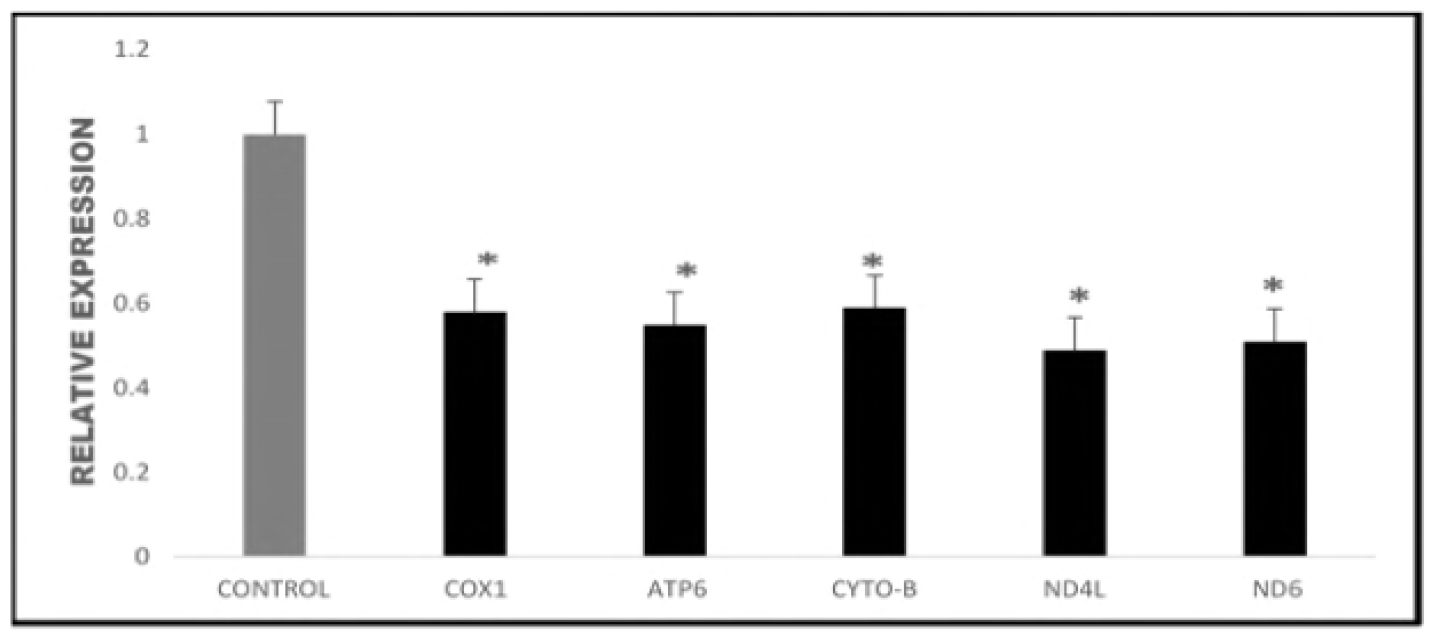
miR 383 down regulates all target genes, mi RNA 383 was overexpressed in HepG2 cells and the mitochondria was isolated. RT-PCR analysis was performed to check the impact of increased miR 383 level on the expression pattern of target mitochondrial genes. The result revealed that the expression of all the mitochondrial target genes of miR 383 went significantly down. Results presented are average of three experiments ± SEM each done at least in triplicate, p<0.05.*Statistically significant when compared to control.

**Figure 4.4:**
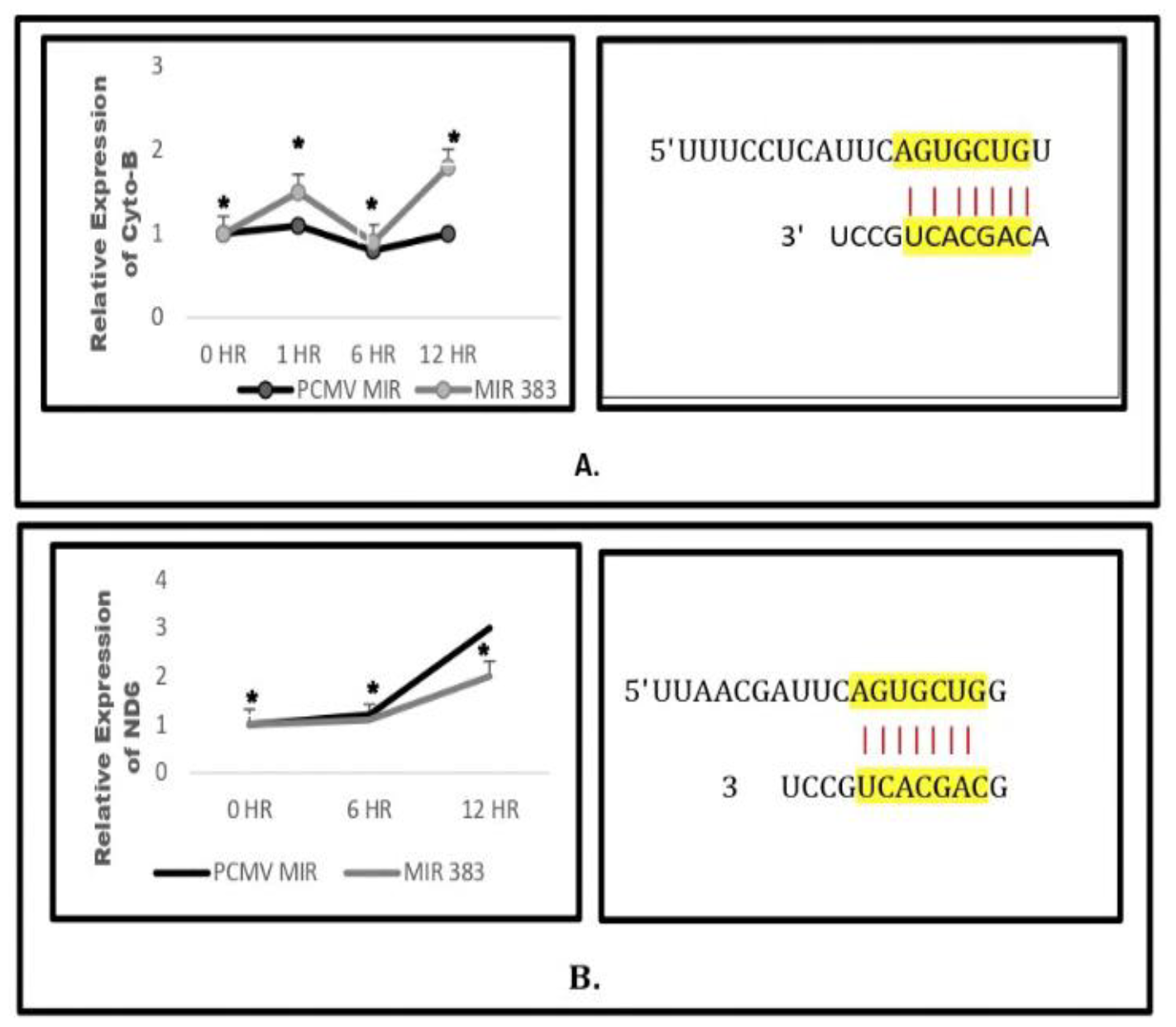
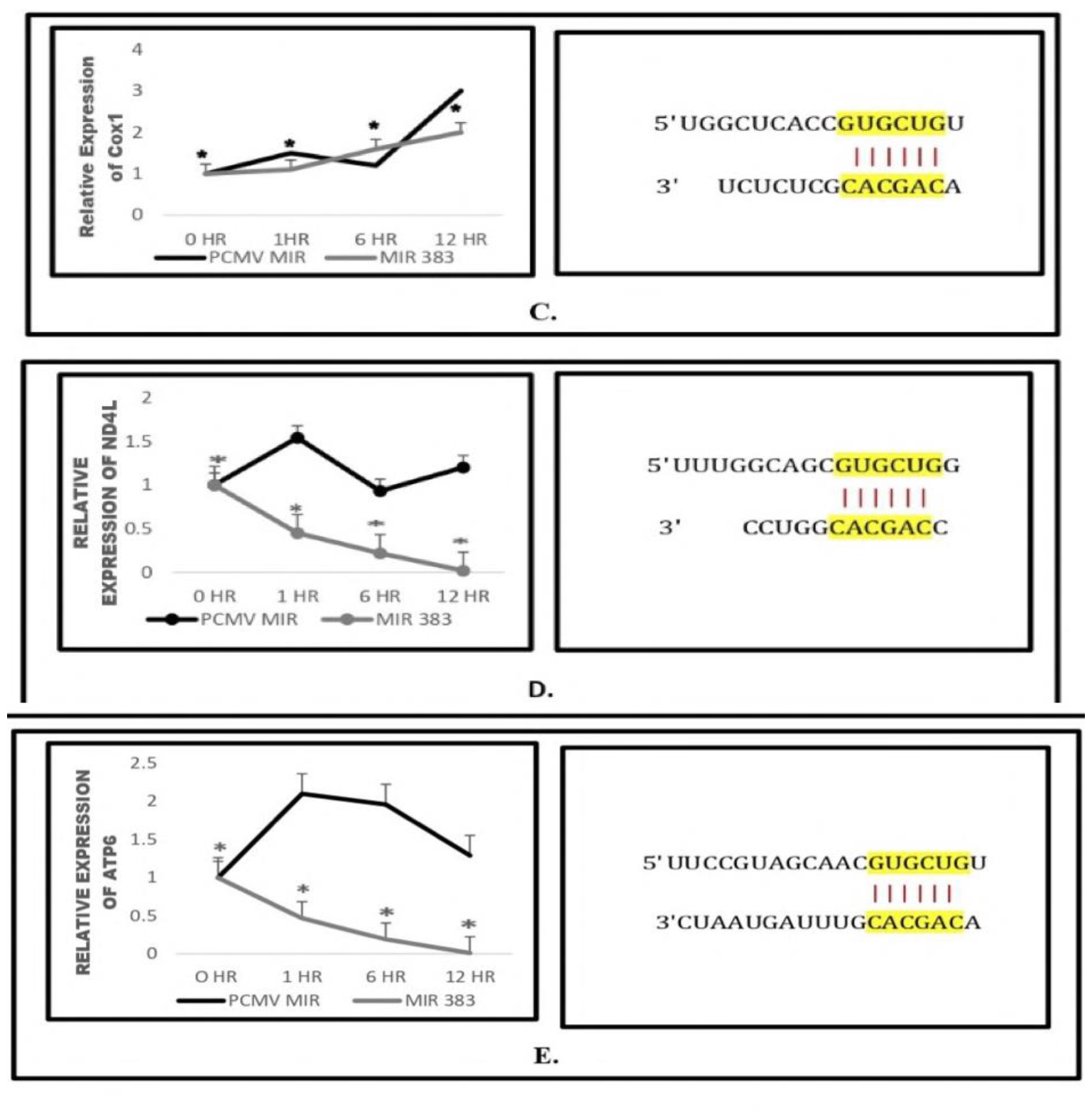
3’ UTR analysis and mRNA stability assay revealed the direct targeting of ATP6 andND4L by miR 383. miR383 wasoverexpressed in Hepg2 cells and 24 hours post transfection, the cells were treated with actinomycin D and RNA samples were collected at 0, 1, 6 and 12 hours respectively. Following this, RT-PCR analysis was performed to check the expression pattern of target genes. Results revealed that expression of ND4L and ATP6 went significantly down with time suggesting its direct targeting by miR383. The expression pattern of Coxl, ND6 and Cyto-B also was altered but not significantly. A.) 3’ UTR analysis and the relative expression of Cyto-b B.) 3’ UTR analysis and the relative expression of ND6C.) 3’ UTR analysis and the relative expression of Coxl D.) 3’ UTR analysis and the relative expression of ND4L E.) 3’ UTR analysis and the relative expression of ATP6. Results presented are average of three experiments ± SEM each done at least in triplicate, P<0.05.*Statistically significant when compared to control.

**Figure 4.5:**
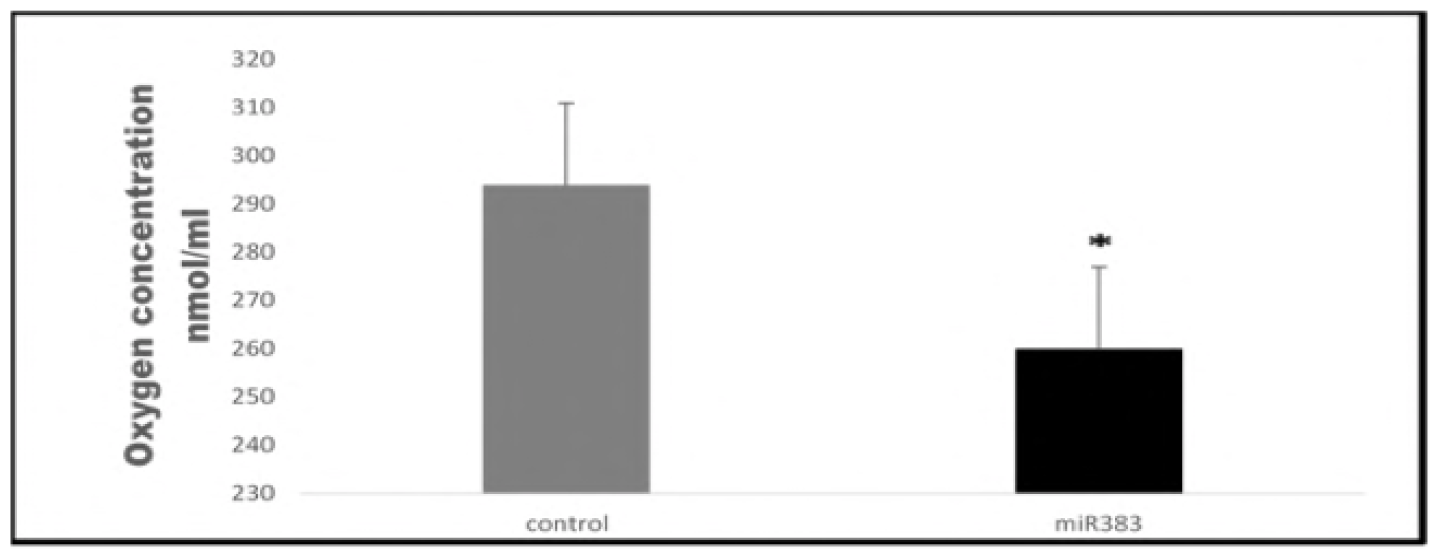
Levels of oxygen consumption by the cells went significantly down, when miR 383 was over expressed. miR 383 was over expressed in HepG2 cell line and 24 hour post transfection, oxygraph analysis was performed. The results revealed a decrease in O_2:_ consumption by the mitochondria of miR 383 overexp res sing cells when compared with the mock transfected cell. Results presented are average of three experiments ± SEM each done at least in triplicate. p< 0.05 .*Statistically significant when compared to control.

**Figure 4.6:**
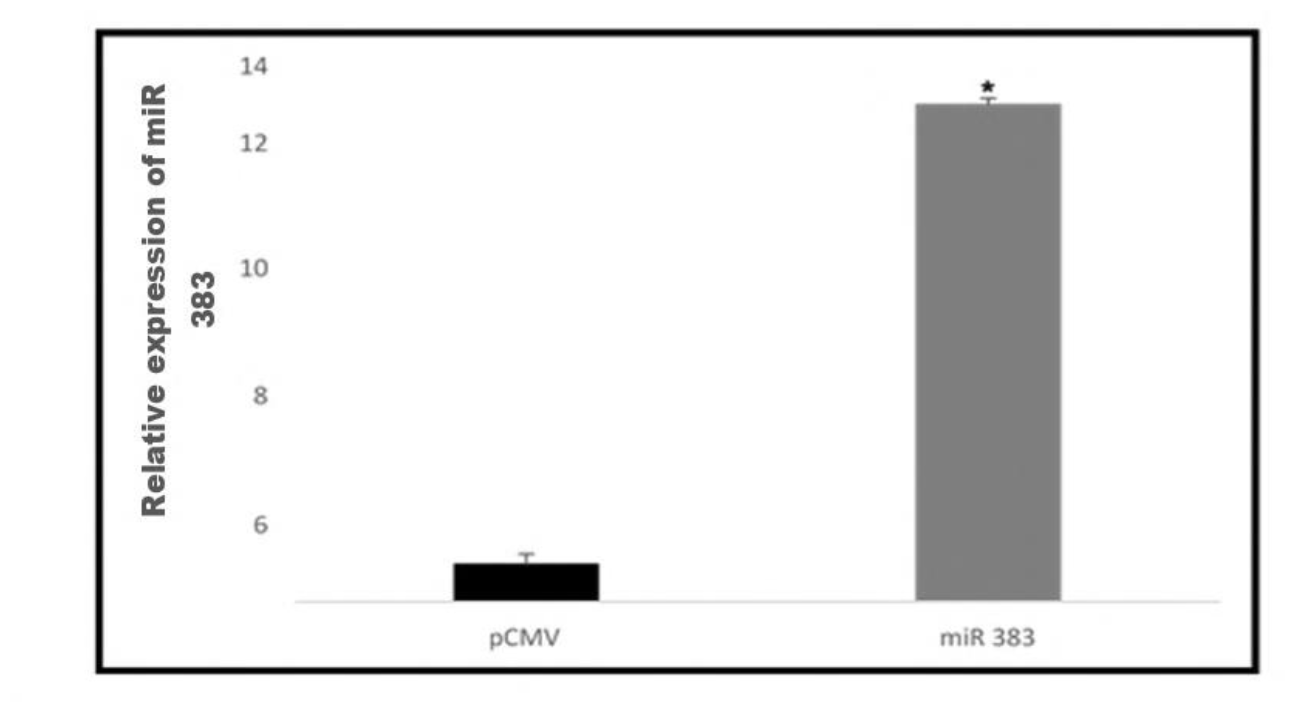
Exosomes isolated from miR 383 over expressing cells were enriched with miR 383. Real time PCR analysis was performed to check the enrichment of miR383 in the exosomes isolated from miR 383 over expressing cells and compared with the exosomes isolated from the control cells. The results revealed that exosomes isolated from microRNA 383 over expressing cells, shown 13 fold higher level of microRNA 383. Results presented arc average of three experiments ± SEM each done at least in triplicate, p<0.05.*Statistically significant when compared to control.

**Figure 4.7:**
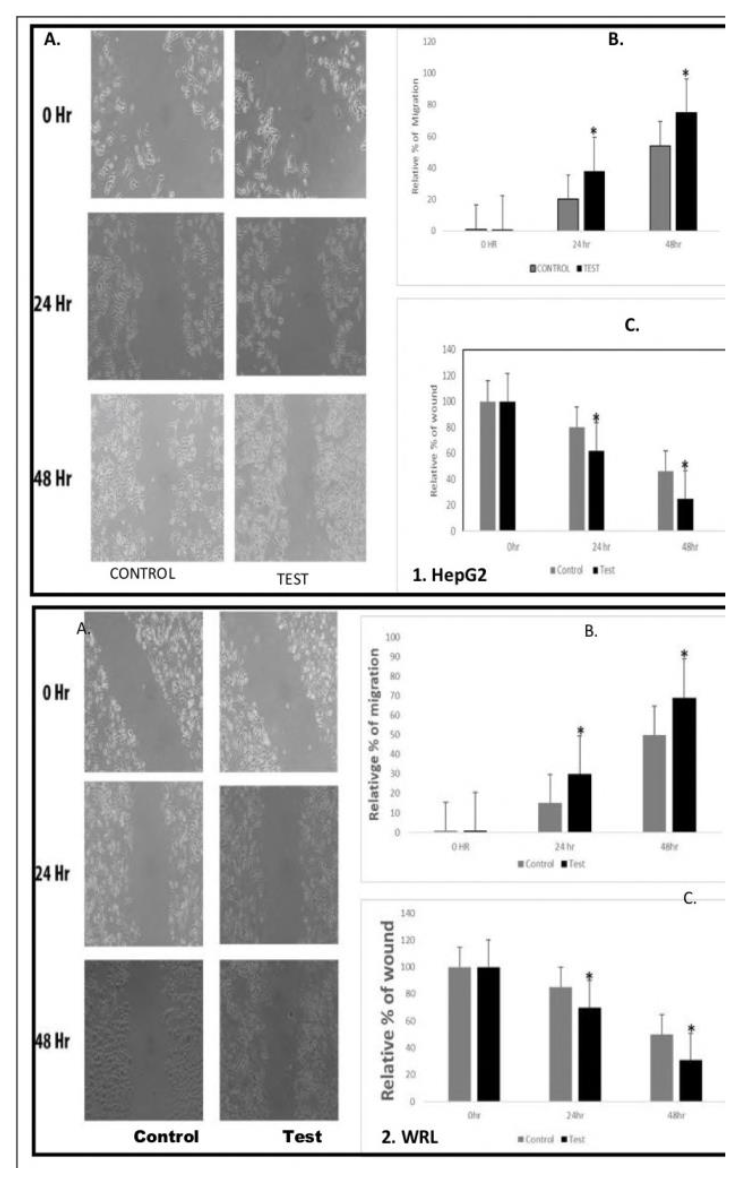
miR 383 escalates the cellular migration when treated with exosomes isolated from microRNA 383 over expressing cells. Cells were grown in a monolayer and a scratch was made, followed by exosome treatment. The microphotographs were taken at 0, 24, and 48 hours. The distance/gap covered by the cells with time was estimated by image-J software. The result revealed that the rate of the cellular migration got escalated when treated with exosomes enriched with miR 383. 1A.) Microphotograph of cell migration pattern (HepG2) with respect to time 1B.) Relative percentage of migration at 0, 24 & 48 hours 1C.) Relative percentage of wound healing at 0, 24 & 48 hours. 2A.) Microphotograph of cell migration pattern (WRL) with respect to time 2B.) Relative percentage of migration at 0, 24 & 48 hours 2C.) Relative percentage of wound healing at 0, 24 & 48 hours. Results presented are average of three experiments ± SEM each done at least in triplicate, p<0.05.*Statistically significant when compared.

**Figure 4.8:**
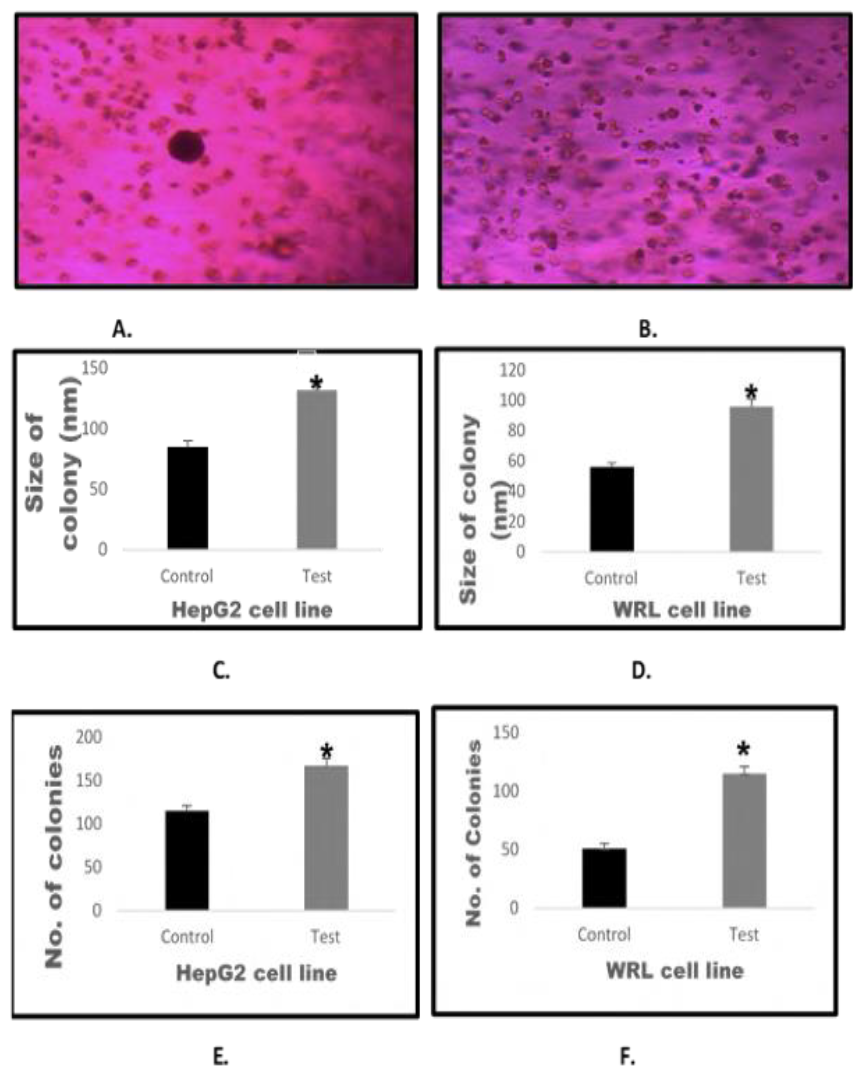
miR 383 enhances the tumor size when treated with exosomes isolated from microRNA 383 over expressing cells. HepG2 and WRL colonies were treated with exo somes iso la ted from the miR 383 overexpressing cells and allowed to grow for 28 days. Following this, the microphotographs were taken at 20 different regions and 20 colonies from each region was taken for the size estimation by image-J software The number of colonies were counted manually from 20 regions to estimate the number of tumor colonies. The results revealed that the size as well as the number of colonies got enhanced in HepG2 and WRL cells, when treated with exosomes enriched with miR383. A.) Micro photograph of HepG2 cell colony B.) Micro photograph of WRL cell colony C.) Comparative colony sizeofHepG2 cells. D.) Comparative colony size of WRL cells E.) Number of colonies of HepG2cells, when compared with control F) Number of colonies of WRL cells, when compared with control. Results presented are average of three experiments ± SEM each done at least in triplicate p<0.05.*Statistically significant when compared to control.

